# The evolution of genetic bandwagoning

**DOI:** 10.1101/064493

**Authors:** Idan S. Solon

## Abstract

**Background:** In separate literatures, biologists have marshaled theoretical and empirical support for theories that a variant can be selected to (1) induce suicide by moribund or otherwise stressed holders; (2) induce suicide by a holder with low productive or reproductive potential; (3) impose senescence upon a chronologically old holder; and (4) reduce the yield of a holder with low genetic heterozygosity. High stress, moribundity, low productive and reproductive potential, high chronological age, and low genetic heterozygosity all indicate the holder has a relative paucity of advantageous genetic variants in acquiring prey, territory, and mates or surviving predators and parasites. Therefore, an unappreciated commonality between these theories is that a variant can be selected that reduces the fitness of a holder of low genetic quality.

Here, I argue that a variant can be selected that reduces its holder’s fitness in response to not just these indications but also other indications of low genetic quality. The fitness losses induced by the variant in low-quality individuals allow fitness gains for nearby individuals, some of which hold a copy of the variant. The variant gains frequency by hitchhiking along with (“jumping on the bandwagon” of) higher-quality individuals (and their lineages) that have copies of the variant; therefore, it is called a “bandwagoning” variant.

**Questions:** What parameter values (e.g., population quantity, relatedness, heritability of reproductive success) allow natural selection of genetic bandwagoning?

**Features of the model:** The model is an individual-based Moran process. Each individual’s quality value is randomly chosen at birth from a normal distribution that has a mean equal to the quality value of its parent.

**Ranges of parameters:** Total population quantity varied from 50 to 500. Assortment (“relatedness”) in the population varied from .05 to .15. Recorded values for the heritability of reproductive success varied from .024 to .132.

**Conclusions:** Natural selection of genetic bandwagoning can occur even when values for population quantity, relatedness, and heritability of reproductive success are low enough to be in line with reported values for humans and other species. Therefore, genetic bandwagoning theory can explain why indications of an organism’s low genetic quality induce behavior by, or biological processes within, that organism that reduce that organism’s fitness.

## 1. Introduction

In various literatures, biologists have argued that natural selection has shaped a conditional tendency to forfeit resources (e.g., food, territory, mating opportunities) so that they can be utilized by any nearby individual(s). The individuals that utilize them are unaware that the resources were even relinquished; therefore, such forfeitures are not explained by direct or indirect reciprocity (Nowak, 2012). A commonality across these theories is that the forfeiture occurs on a particular condition that indicates the individual is lower-quality^1^ compared to its neighbors. However, biologists advancing these theories have seldom explicitly stated a condition of low genetic quality.

For example, multiple biologists have hypothesized that if an individual is moribund due to a parasite infection or some other malady, it might commit suicide, thereby relinquishing the individual’s resources to nearby individuals, whether the individuals benefiting are kin (e.g., Dawkins, 1976) or not (Refardt et al., 2013). Additionally, some biologists have argued that programmed cell death occurs in unicellular organisms such as *Escherichia coli, Caenorhabditis elegans*, amoeba, and yeast on the condition that the cell is stressed, even if the stress itself does not suggest imminent death (Skulachev, V., 1999, 2001, 2002; Thompson and Kay, 2000; Hazan et al., 2004; Herker et al., 2004; Pestov et al., 2011). Individuals that have less advantageous heritable traits compared to neighbors are more likely to incur moribundity and various other stressors.

Furthermore, for de Catanzaro (1981, 1984), depression and its associated suicidal ideations occur in individuals with low reproductive and productive potential in order to leave resources to more productive kin. He argued (1984, pg. 77) that “the death of individuals with seriously impaired reproductive and productive potential might actually benefit their inclusive fitness by conserving resources for kin not experiencing such impediments.” Low reproductive and productive potential can have a genetic basis, in which case the depression would be incurred by low-quality individuals.

Other biologists have argued that natural selection can shape a tendency to forego resources even if the foregoing individual is not moribund or closely related to neighboring individuals. For example, recent years have seen a resurgence of Weismann’s (1889) old theory that natural selection has occurred for senescence to occur in conjunction with chronological age, i.e., programmed ageing (Mitteldorf, 2004, 2006; Longo et al., 2005; Goldsmith, 2008, 2014; Pepper et al., 2013; Skulachev, M. & Skulachev, V., 2014; Mitteldorf & Sagan, 2016). Some authors hold that programmed ageing occurs because aged individuals can be expected, due to their chronological age, to be of low *genetic* quality (Skulachev, V., 1997; Goldsmith, 2004; Travis, 2004; Martins, 2011; Yang, 2013).

Additionally, Semel et al. (2006) noted that, in populations of tomato, rice, and maize, the overdominance associated with heterosis—the higher fitness of hybrids compared to their inbred parents—occurs primarily among reproductive traits. They argued this constitutes evidence for natural selection of heterosis itself—that is, natural selection in favor of the association between hybrids and higher fitness—in order to promote heterozygosity in populations in a manner analogous to a plant’s self-incompatibility system. The implication is that there has been natural selection in favor of this association between outbred individuals and yield improvements because heterozygous individuals are of higher genetic quality.

Here, I introduce the term genetic bandwagoning^2^ to refer to a variant that induces the individual in which it is located to forfeit some or all^3^ of the resources (e.g., food, territory, mates) it could have used if that individual’s quality is sufficiently low compared to neighbors. The heritable elements (e.g., alleles, epigenetic marks) within a single individual that are responsible for bandwagoning are collectively considered a “bandwagoning variant.”

The bandwagoning variant would not know which neighbors would hold a copy of the same variant. Some resources would be used by individuals that do not. However, if the difference in quality between the forgoing individual and the neighbors is sufficiently substantial, neighbors that hold a copy of the variant can gain more descendants by using a fraction of the resources and reproductive opportunities than the forgoing individual could by using all. The reason is that descendants of the higher-quality individuals are likely to be better at surviving and reproducing and, therefore, have a higher fitness per descendant than the descendants of the foregoing individual.

Genetic bandwagoning is an example of the Hankshaw effect—by which “a property of an allele increases its likelihood of hitchhiking” (Hammarlund et al., 2016, pg. 1376)—because when copies of a bandwagoning variant induce low-quality individuals to forfeit resources, higher-quality individuals gain resources, facilitating hitchhiking by copies of the variant held by higher-quality individuals. Moreover, this hitchhiking capacity is less likely to be reduced by sex and recombination than some forms of hitchhiking, since the bandwagoning variant hitchhikes with not just one allele but a multiplicity of alleles and epigenetic marks responsible for an individual’s high quality. Consequently, after sex and recombination, the variant is likely to continue to be located with half of the alleles to which the individual’s high quality was originally attributable and the parent that contributes the other half of the alleles with which the variant is located is also likely to be high-quality, insofar as there is assortative mating by quality as a result of sexual selection by both sexes (Bos et al., 2009; Griggio and Hoi, 2010; Holveck and Riebel, 2010; Holveck et al., 2011; Dakin and Montgomerie, 2014; Veen and Otto, 2015; Schultzhaus et al., 2017).

Consider that a low-quality actor would produce *C_1_* offspring if it does not relinquish resources. Alternatively, if the actor relinquishes resources, they are used by *n* recipients; the *i*^th^ recipient gains *B*_*i*1_ offspring and is related to the actor by *r*_i_. According to Hamilton’s (1964) rule, a strategy of relinquishing resources on the condition of low quality—that is, genetic bandwagoning—can gain frequency by natural selection after one generation if:

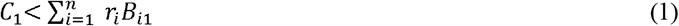

The foregoing inequality is consistent with most analyses of the evolution of social behavior, which examine solely the costs and benefits that result from an act by the first generation following the act (Hunt et al., 2004). However, this underestimates the likelihood that genetic bandwagoning evolves. Not only do higher-quality individuals produce, on average, more offspring, but these offspring are also likely to be, on average, higher-quality, since genetic quality is, by definition, heritable. Historically, theory has predicted variation in genetic quality to be low (Borgia, 1979; Taylor and Williams, 1982). However, in recent decades, it has become accepted that mutations (Pomiankowski and Moller, 1995; Rowe and Houle, 1996) and epigenetic changes (Bonilla et al., 2016) can maintain variation in genetic quality.

The offspring of the low-quality actor and their later descendants in the following generations may also be, on average, low-quality themselves compared to the offspring and later descendants of the recipients. The strategy of genetic bandwagoning can gain frequency by natural selection by generation *v* if:

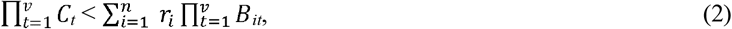

where *C_t_* is the quantity of offspring produced, on average, by the descendants of the low-quality actor during generation *t* and *B_it_* is the quantity of offspring produced, on average, during the same generation *t* by the descendants of the *i^th^* recipient.

The bandwagoning variant can evolve by natural selection in the same way that a person that pays an expert investor to invest his capital can make more money overall over a number of years than if he invests his own capital—even if he has to pay the investor a high percentage of his capital up front—because the expert investor is better at making money. The evolution of the bandwagoning variant owes to the higher fitness per descendant of the higher-quality individuals, just like the success of the investment owes to the higher percentage per year earned by the expert investor.

## 2. Model

The model used herein is a Moran (1958) process, which is chosen for its simplicity and prevalence of use in evolutionary theory (Lieberman et al., 2005; Nowak, 2006a; Proulx, 2011; Shakarian et al., 2012). The Moran model’s fundamental characteristic is that during each time-step, one individual in the population is randomly chosen to produce a single offspring and one individual is randomly chosen for death. (It is possible for the individual that reproduces in a time-step to also be the individual that perishes in the same time-step.) Therefore, the population’s size remains a constant size *N* from one time-step to another.

**Table 1.** Baseline parameter^4^ values

| Symbol | Description | Value |
| --- | --- | --- |
| $N$ | population size | 200 |
| $N_C$ | starting quantity of the cooperator type | 100 |
| $N_D$ | starting quantity of the defector type | 100 |
| $Q_\sigma$ | standard deviation of distribution from which an individual's quality is generated | 1 |
| $Y$ | fraction of fitness lost that is used by other individuals in the population | 0.8 |
| $R$ | the assortment (“relatedness”) in the population | 0.15 |
| $P$ | quality percentile cutoff for reproduction | 10 |
| $T$ | number of time-steps in each run | 5000 |

<sup>4</sup> Some of the model's baseline values are discussed in Appendix section A2 and some simplifications included in the model are discussed in Appendix section A3.

Each individual, *i*, in the population is characterized by a number, denoted *Q_i_*, that represents that individual’s quality. During the initial time-step, the quality value, *Q_i_*, of each of the *N* individuals in the population is randomly generated from a normal distribution with a mean equal to 10 and a standard deviation of *Q_σ_*. (The random generation of quality values for individuals born after the first time-step is described in section 2.2.)

There are two types of individuals in the population: cooperators and defectors. Cooperators are of the bandwagoning type, so a cooperator relinquishes its opportunity to reproduce in a time-step if its quality is below the *P* percentile of all individuals (regardless of type) in the population. Defectors do not do bandwagoning.

### 2.1 Other individuals benefit from fitness foregone by low-quality cooperators

An individual, *i*, in the population (whether the individual is a cooperator or defector) has fitness that is equal to that individual’s quality, *Q_i_*, plus its share of fitness foregone by cooperators with quality below the *P* percentile. (A cooperator with quality below the *P* percentile has fitness of zero irrespective of its quality and its fitness foregone is equal to that individual’s quality.)

A higher share of this foregone fitness goes to other cooperators due to an assumption of positive assortment. Additionally, other individuals in the population gain a share of foregone fitness that is in proportion to their quality. (Precise calculations of fitness are given in Appendix section A1.)

### 2.2 Birth and death

In each time-step, one individual in the population is randomly chosen for reproduction and one individual in the population is randomly chosen for death. (The same individual may be chosen for both reproduction and death.) The probability of being chosen for reproduction is proportional to fitness. The individual chosen for death is determined by randomly choosing N – 1 individuals for survival. The individual not chosen for survival is the one that perishes. The probability of being chosen for survival is proportional to quality.

Once an individual is chosen for reproduction, an offspring of the same type is born and replaces the perished individual. The quality value of its offspring is randomly selected from a normal distribution with a mean equal to its parent’s quality and a standard deviation of *Q_σ_* (the same standard deviation of the normal distribution used to generate the quality values of the *N* individuals at the beginning of the simulation). This random generation of an offspring’s value for quality models the effect of germline mutation, as the quality values for offspring vary randomly about their parent’s value. Since an offspring’s quality value is drawn randomly from a normal distribution centered at its parent’s value, it can potentially be either lower than, equal to, or higher than its parent’s value, which would represent, respectively, a negative, neutral (Kimura, 1968), or positive (Waite and Shou, 2012) net influence of mutations upon quality. Quality is, therefore, partly but not completely heritable.

After the birth of one individual and the death of one individual, the quality value of each individual in the population is multiplied by a normalizing factor^5^, 10*N*/*Q_T_*, where *Q_T_* is the sum of the quality values of all the individuals in the population. The run continues for *T* time-steps or until one type has gained fixation.

## 3. Results

Results are graphically represented in Figs. 1, 2, and 3. A pattern common to these figures is the drop, on average, of cooperator quantity early in runs as the bandwagoning variant essentially sacrifices holder quantity for holder quality. This occurs as low-quality cooperators forego reproduction, which enhances the fitness of other individuals in the population. The fitness gained by other cooperators from this foregone reproduction is less than the fitness lost from it by low-quality cooperators. Therefore, cooperator quantity is likely to be lower in the following time-step than if lower-quality cooperators had not foregone reproduction. However, since assortment is positive, it is likely that each higher-quality cooperator gains more fitness from the foregone reproduction than each defector with similarly high quality. Therefore, the likelihood that a cooperator produces a high-quality offspring increases compared to the likelihood that a defector produces a high-quality offspring. A higher number of high-quality offspring for cooperators leads to cooperator quantity gains later in runs, as high-quality individuals tend to have more descendants.

**Fig. 1.**
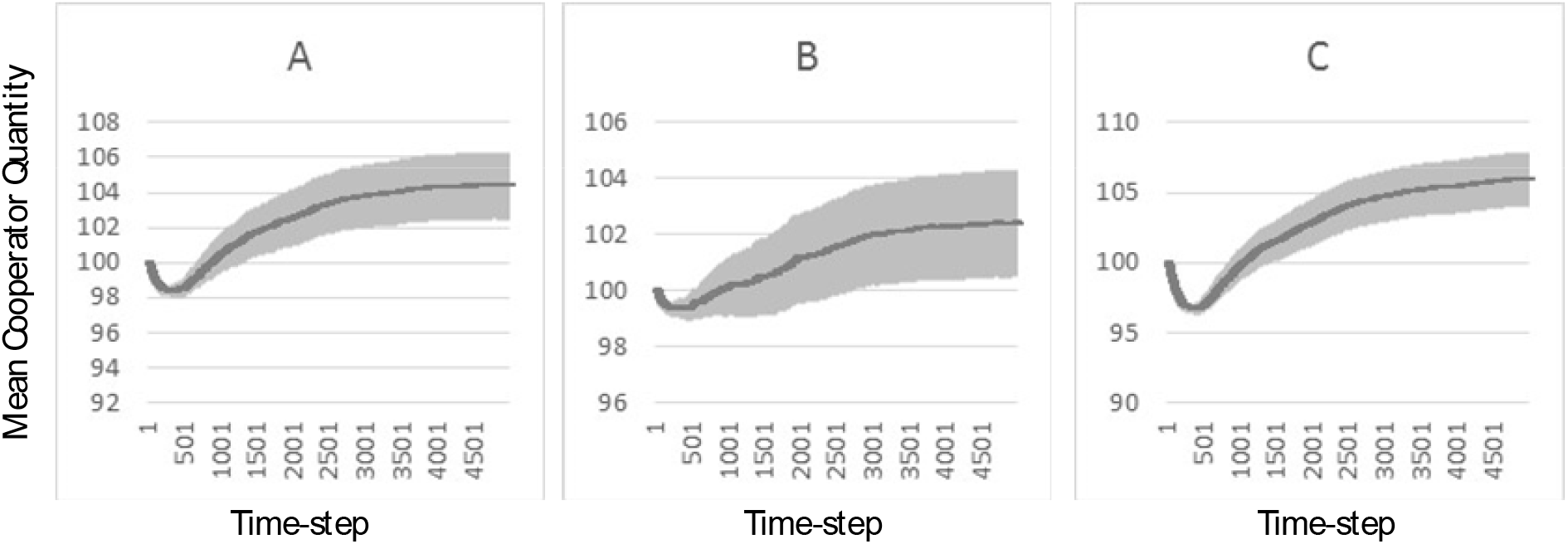
The average quantity of cooperators across 10000 replicate runs is indicated by the dark line. Shaded regions indicate 95% confidence intervals. For purposes of calculating these averages and confidence intervals, when defectors gained fixation during a run, then cooperator quantity was tallied as zero for each time-step thereafter for that particular run and when cooperators gained fixation during a run, then cooperator quantity was tallied as *N* for each time-step thereafter for that particular run. Heritability (*h^2^*) values for reproductive success (RS) were calculated using the values recorded as the program ran. The reported *h^2^* RS values are the per-run values averaged across 10000 runs. (A) With all values equal to the baseline values in Table 1, the cooperator quantity was, on average, significantly above the starting cooperator quantity of 100 by the 5000^th^ time-step. *h^2^* RS = .086. (B) With all values equal to the baseline values except *P* = 5, the cooperator quantity was, on average, significantly above the starting cooperator quantity of 100 by the 5000^th^ time-step. *h^2^* RS = .085. (C) With all values equal to the baseline values except *P* = 15, the cooperator quantity was, on average, significantly above the starting cooperator quantity of 100 by the 5000^th^ time-step. *h^2^* RS = .087.

**Fig. 2.**
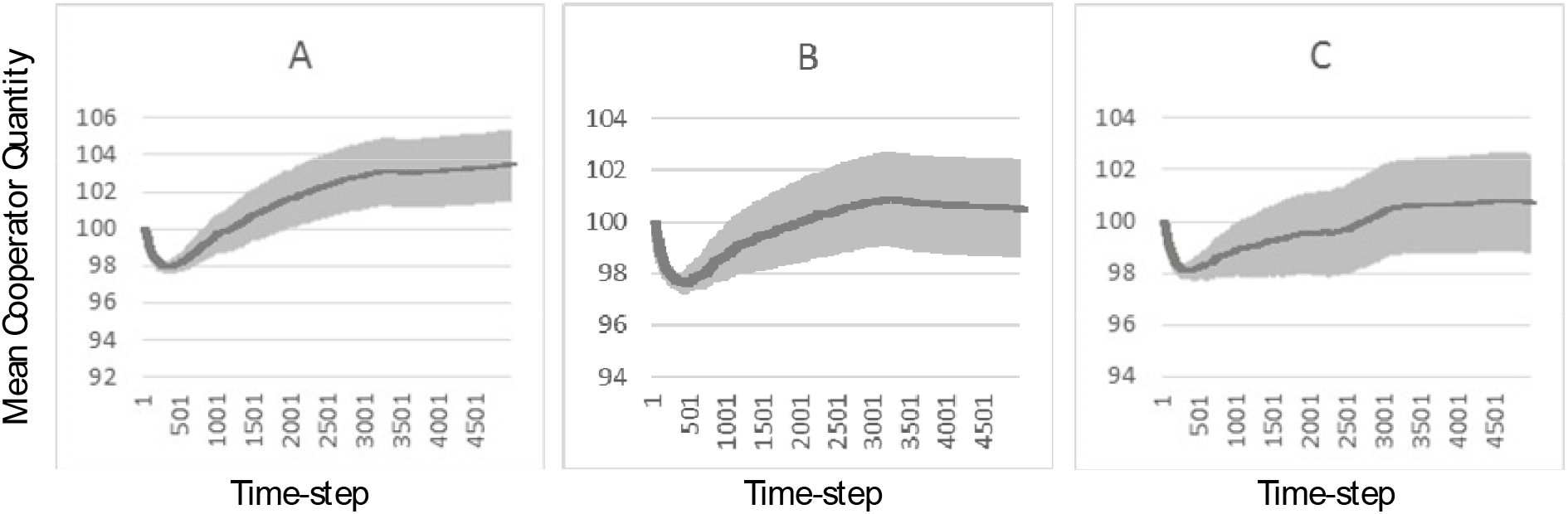
The average quantity of cooperators across 10000 replicate runs is indicated by the dark line. Shaded regions indicate 95% confidence intervals. For purposes of calculating these averages and confidence intervals, when defectors gained fixation during a run, then cooperator quantity was tallied as zero for each time-step thereafter for that particular run and when cooperators gained fixation during a run, then cooperator quantity was tallied as *N* for each time-step thereafter for that particular run. Heritability (*h^2^*) values for reproductive success (RS) were calculated using the values recorded as the program ran. The reported *h^2^* RS values are the per-run values averaged across 10000 runs. (A) With all values equal to baseline values except r = .1, the cooperator quantity was, on average, significantly above the starting cooperator quantity of 100 by the 5000^th^ time-step. *h^2^* RS = .086. (B) With all values equal to baseline values except *r* = .05, the cooperator quantity was, on average, higher than the starting cooperator quantity of 100 by the 5000^th^ time-step, but the margin was not significant. *h^2^* RS = .086. (C) With all values equal to baseline values except *Y* = .5, the cooperator quantity was, on average, higher than the starting cooperator quantity of 100 by the 5000^th^ time-step, but the margin was not significant. *h^2^* RS = .086.

**Fig. 3.**
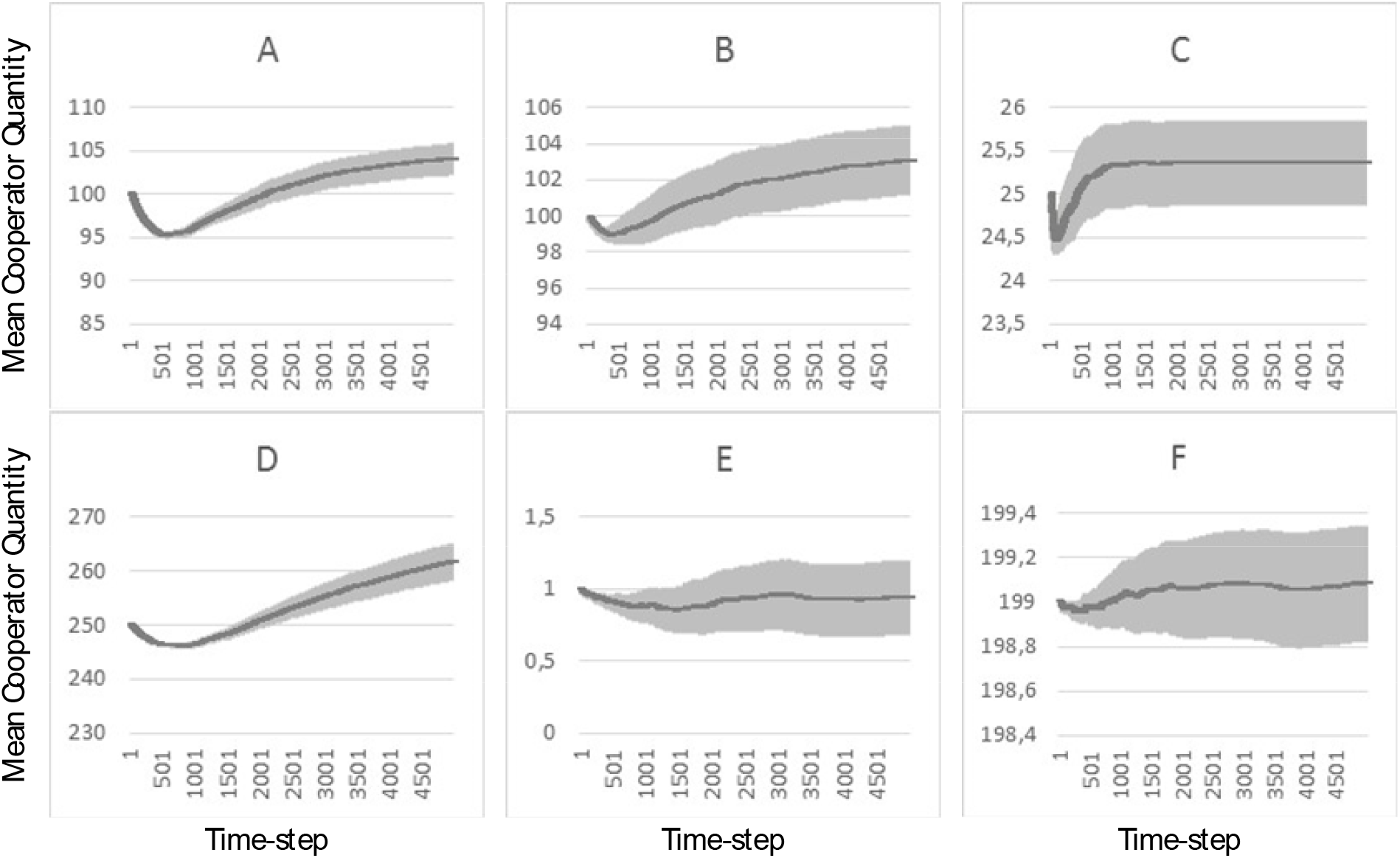
The average quantity of cooperators across 10000 replicate runs is indicated by the dark line. Shaded regions indicate 95% confidence intervals. For purposes of calculating these averages and confidence intervals, when defectors gained fixation during a run, then cooperator quantity was tallied as zero for each time-step thereafter for that particular run and when cooperators gained fixation during a run, then cooperator quantity was tallied as *N* for each time-step thereafter for that particular run. Heritability (*h^2^*) values for reproductive success (RS) were calculated using the values recorded as the program ran. The reported *h^2^* RS values are the per-run values averaged across 10000 runs. (A) With all values equal to baseline values except *Q_σ_* = .3, the cooperator quantity was, on average, significantly above the starting cooperator quantity of 100 by the 5000^th^ time-step. *h^2^* RS = .032. (B) With all values equal to baseline values except *Q_σ_* =3, the cooperator quantity was, on average, significantly above the starting cooperator quantity of 100 by the 5000^th^ time-step. *h^2^* RS = .132. (C) With all values equal to baseline values except N_C_ = N_D_ = 25, the cooperator quantity was, on average, above the starting cooperator quantity of 25 by the 5000^th^ time-step, but the margin was not significant. *h^2^* RS = .053. (D) With all values equal to baseline values except N_C_ = N_D_ = 250, the cooperator quantity was, on average, significantly above the starting cooperator quantity of 250 by the 5000^th^ time-step. *h^2^* RS = .103. (E) With N_C_ = 1; N_D_ = 199; and all other values equal to baseline values; mean cooperator quantity was, on average, lower than the beginning cooperator quantity of 1 by the 5000^th^ time-step, but the margin was not significant. *h^2^* RS = .024. The cooperator type gained fixation in 47 of 10000 runs. (F) With N_D_ = 1; N_C_ = 199; and all other values equal to baseline values; mean cooperator quantity was, on average, higher than the beginning cooperator quantity of 199 by the 5000^th^ time-step, but the margin was not significant. *h^2^* RS = .027. The defector type gained fixation in 46 of 10000 runs.

With variables equal to baseline values, cooperator quantity, on average, exceeded starting cooperator quantity of 100 later in runs (Fig. 1A). This result was relatively robust to changes in the quality percentile cutoff for reproduction. As Figs. 1B and 1C show, when cooperators reproduced only if their quality percentile was, respectively, at least 5 and at least 15, cooperator quantity, on average, exceeded starting cooperator quantity later in runs. In all three cases, the margin by which mean cooperator quantity exceeded 100 later in runs was significant. This demonstrates that the success of a bandwagoning strategy does not require an individual to identify with absolute precision whether its quality is below a particular percentile.

For Figs. 2A to 2C, one variable value was changed to be less favorable to cooperators and all other variables were equal to baseline values. With *r* = .1 (Fig. 2A), cooperator quantity, on average, significantly exceeded starting quantity of 100 later in runs. However, the margin by which cooperator quantity, on average, exceeded starting quantity of 100 later in runs was not significant when *r* = .05 (Fig. 2B) or *Y* = .5 (Fig. 2C).

For Figs. 3A and 3B, respectively, higher and lower values were used for the standard deviation of the normal distributions from which offspring quality values were generated. All other variable values were equal to baseline. In both figures, cooperator quantity, on average, significantly exceeded beginning quantity of 100 later in runs. For Figs. 3C and 3D, respectively, lower and higher total population quantities were used. All other variable values were equal to baseline. In Fig. 3C, cooperator quantity, on average, exceeded beginning cooperator quantity of 25 later in runs, but the margin was not significant. In Fig. 3D, cooperator quantity, on average, exceeded beginning cooperator quantity of 250 later in runs and the margin was significant. A higher total population quantity probably tended to benefit cooperators because a higher total population quantity made it less likely that defectors gained fixation when the percentage of cooperators dipped early in runs, enabling cooperator percentage to recover later in runs.

The results also demonstrated that the cooperator type can invade the defector type. For Fig. 3E, cooperator quantity started at 1, defector quantity started at 199, and other variable values were equal to baseline values. The cooperator type gained fixation in 47 of 10000 runs. For Fig. 3F, cooperator quantity started at 199, defector quantity started at 1, and other variable values were equal to baseline values. The defector type gained fixation in 46 of 10000 runs.

## 4. The evaluation of a holder’s quality

The assumption that a variant can make a useful approximation of an organism’s quality pervades evolutionary theory. For example, both the “sexy son” (Fisher, 1930; Weatherhead and Robertson, 1981; Andersson, 1994) and “good genes” (Andersson, 1982, 1994; Iwasa and Pomiankowski, 1994; Moller and Alatalo, 1999; Byers and Waits, 2006) hypotheses of sexual selection involve an assessment by the female about the reproductive value of a male’s offspring (Kokko et al., 2002). Additionally, parents in many species are believed to identify which of their offspring have the best genes for the purposes of allocating more resources to those offspring (Burley, 1986; Sheldon, 2000; Harris, W. and Uller, 2009). Moreover, parents are theorized to kill low-quality and/or less attractive offspring by practicing filial cannibalism (Klug and Bonsall, 2007) and, more generally, parental selection (Harris, J., 2006).

In the above contexts, the organism whose quality is assessed and the organism making the assessment have competing interests, due to sexual conflict (Williams, 1966; Parker, 1979) and/or parent-offspring conflict (Trivers, 1974). A misalignment of the interests of signaler and receiver is a factor that encourages signaling dishonesty (Maynard Smith, 1991; Hurd and Enquist, 2005; Searcy and Nowicki, 2005; Szamado, 2011). However, prevailing theory holds that such assessments can be made accurately, primarily on the basis of honest signals of quality (Andersson, 1994; Nowicki et al., 1998; von Schantz et al., 1999; Duffield et al., 2017). By contrast, a variant’s bandwagoning activity is compatible with the interests of the variants at other loci of the same individual. That is, the values that apply to the variables in inequality (3) that determine whether a variant can gain frequency by natural selection from doing bandwagoning are the same as the values that apply to determining whether another variant within the same individual can gain frequency through the same bandwagoning, even if that other variant is not responsible for bandwagoning. This suggests information (e.g., hormonal, physiological) received by the variant about its holder’s quality is at least as reliable as information about an organism’s quality that is received through honest signals of quality by potential mates or parents of that organism. Yet bandwagoning theory does not require a variant that evaluates its holder’s quality to have any particular foresight or to make an evaluation with any more precision than variants are assumed to have when they evaluate another organism’s quality.

Indeed, theory holds that organisms can approximately evaluate the fitness they would obtain for the purpose of practicing fitness-dependent sex (Hadany and Beker, 2003; Hadany and Otto, 2007; Ram and Hadany, 2016). Empirical support has come so far from studies of microbes, fungi, and plants, but the theory is thought to be applicable to other species (Ram and Hadany, 2016).

Information that allows an estimation about the holder’s quality can potentially come from a variety of sources. Honest signals indicate quality because they are diminished when organisms experience various stressors, such as hunger, infection, thermal, or psychosocial stressors (Andersson, 1994; Nowicki et al., 1998; von Schantz et al., 1999; Duffield et al., 2017). A bandwagoning variant could conceivably use biochemical, endocrinological, and physiological indications of such stressors (Pickering and Pottinger, 1995; Sapolsky, 2001; Schneiderman et al., 2005) as well. Additionally, theory holds that an organism is aware of its position in a dominance hierarchy and that this position is partly determined by the organism’s quality (Hsu et al., 2006; Georgiev et al., 2015).

Other phenomena noted above—mate choice, differential allocation, and parental selection—are also potential indicators to an organism of that organism’s quality. Indeed, when parents and prospective mates give to an organism more or less resources or attention, they wind up communicating information to that organism about how its quality compares with conspecifics.

Additionally, in some phenomena that are offered (section 1) as examples of bandwagoning, a bandwagoning variant uses an organism’s older age or low genetic heterozygosity to indicate that the organism is of lower quality.

## 5. A variant is less likely to commit suicide if its quality and lineage fitness outlook are likely to change

As the foregoing suggests, a variant’s evaluation of its holder’s quality and lineage fitness outlook is developed continuously as more and more information becomes available in the form of, for example, different types of stressors; competitive bouts; and interest from potential mates. Such an evaluation is subject to change as new information becomes available. The likelihood that an individual’s lineage fitness outlook changes may depend upon a number of factors that are specific to species, population, and context.

Whether a bandwagoning variant might evolve to induce a holder with a low lineage fitness outlook to relinquish some or all of the holder’s resources would probably depend upon how likely that organism’s lineage fitness outlook is to change in the future. For example, other individuals that are higher-quality might wind up perishing, while the holder survives due to luck. Or the holder’s quality might change because a natural enemy population evolves so that the holder becomes more or less susceptible to that natural enemy population compared to conspecifics. Or the holder’s offspring might be born with a favorable mutation, while the offspring of other organisms are not as lucky. If the holder’s lineage fitness outlook is unlikely to change, it is more likely that a bandwagoning variant would induce the holder to forfeit all of the resources potentially available to the holder (i.e., induce its holder to commit suicide). If the holder’s low lineage fitness outlook is more likely to change, then it is more likely that a bandwagoning variant would induce the holder to forfeit only some resources so that the holder can remain alive in case its fitness outlook changes. This partial forfeiture of resources could involve resource forfeitures over multiple time steps, each contingent upon the holder’s continued low lineage fitness outlook. However, in the model given in section 2, for simplicity, a bandwagoning variant’s decision to induce relinquishment of resources or not occurs only once.

## 6. Conclusions

The thesis of this manuscript is that blind altruism (altruism done anonymously and without preference to related individuals) can evolve by natural selection, provided that the altruism occurs on the condition that the altruistic individual is low-quality. A variant (allele, epigenetic mark, or some combination thereof) responsible for this altruism is, herein, called a bandwagoning variant. The altruism would temporarily reduce the quantity of bandwagoning variant holders, but figures, if assortment is sufficiently positive, to lead to more offspring produced by the higher-quality holders than offspring produced by higher-quality individuals that do not hold the bandwagoning variant. This increases the likelihood that the highest-quality offspring in the population hold a bandwagoning variant, which increases the likelihood that the bandwagoning variant will gain fixation. It boils down to a short-term sacrifice of holder quantity for holder quality.

The model (section 2) was based upon assumptions that are generally supported by prevailing evolutionary theory. The model was developed somewhat conservatively. That is, some of the model’s simplifications were unfavorable to bandwagoning. (The model’s simplifications are discussed in Appendix section A3.) The results (section 3) demonstrated that bandwagoning can evolve when parameter values are in line with those found in the natural populations of some species (see Appendix section A2 about how baseline variable values were chosen).

Furthermore, it is documented in section 1 that a number of scholars have advanced support for theories that altruism that is done anonymously and that is not directed to kin can still evolve by natural selection if it is done on the condition that the altruist is characterized by high stress, high chronological age, low reproductive or productive potential, or low heterozygosity. These characteristics are all indications of low quality, so the theoretical and empirical support these scholars have marshalled for their theories add increasing credibility to the notion that in populations for numerous species, parameters are suitable for the evolution of genetic bandwagoning.

Several theories reviewed in section 1 can be re-interpreted as special cases of bandwagoning theory: the theory of heterosis and the theories holding that programmed ageing evolved because aged individuals can be expected to be of low genetic quality. Evidence that bandwagoning can evolve by natural selection in the specific forms of programmed ageing and heterosis lends credibility to the theory that bandwagoning in whatever form can evolve by natural selection.

Support for the theories of stress-induced suicide and depression reviewed in section 1 is also consistent with bandwagoning theory, but these do not constitute special cases of bandwagoning theory because those theories do not state that the forfeiture is contingent upon low genetic quality. In a departure from de Catanzaro’s theory of depression (see section 1), bandwagoning theory does not require that relinquished resources go to kin, which allows a response to a classic rejoinder to de Catanzaro’s theory of depression: “why wouldn’t a burdensome individual simply leave their kin” instead of committing suicide (Syme et al., 2016, pg. 189)? The answer is that even if relatedness between the relinquishing individual and those benefiting is equal to (or lower than) the relatedness among human populations estimated by Harpending (2002), the benefit-to-cost ratio from suicide can be sufficiently high to satisfy Hamilton’s (1964) rule (per the results in section 3). Therefore, even if a person with low quality were to venture away from his or her family to a different location in the same population, that individual’s forfeiture of resources at that location would still be likely to satisfy Hamilton’s (1964) rule.

Additionally, these examples serve to show that the indications of an organism’s low quality and the ways in which a low-quality organism can be altruistic can take a variety of forms. For instance, low quality can be indicated by high age, low heterozygosity, high stress, or other factors. The altruism can involve suicide or not. As discussed in section 5, the more likely it is for a low-quality holder’s lineage fitness outlook to change, the less likely that holder would be to commit suicide and the more likely it would be to relinquish some resources while also remaining alive in case its lineage fitness outlook changes.

This serves to illustrate that a range of paradoxical empirical phenomena can potentially be explained in this manner. These phenomena would have in common the characteristic that an indication of an organism’s low quality and/or low lineage fitness outlook induces behavior by, or biological processes within, that organism that appear to reduce the fitness of that organism.

### 6.1 Genetic bandwagoning as a source of the evolution of cooperation

The evolution of cooperation is a perennial question (Hamilton, 1964; Trivers, 1971; Nowak, 2006b, 2012; West et al., 2007; West et al., 2011). Cooperation is defined (West et al., 2007) as “a behaviour which provides a benefit to another individual (recipient), and which is selected for because of its beneficial effect on the recipient.” The resource forfeitures by low-quality individuals that result from genetic bandwagoning constitute cooperation that affords indirect genetic benefits (West et al., 2007; West et al., 2011) to the forfeiting individual. In comprehensive reviews of the evolution of cooperation, kin discrimination, green-beard discrimination, group selection, limited dispersal, and spatial selection have been offered as ways by which cooperation for indirect benefits can evolve (Nowak, 2006b, 2012; West et al., 2007; West et al., 2011). These are all ways of referring to a state in which cooperation satisfies Hamilton’s rule primarily via high relatedness between the cooperating and benefiting individuals, rather than a high benefit-to-cost ratio associated with the cooperation. Genetic bandwagoning is distinct from these explanations because cooperation that owes to genetic bandwagoning can satisfy Hamilton’s rule even if relatedness is low, since it satisfies Hamilton’s rule (which is *C* < *rB*) primarily via a high benefit-to-cost ratio (high *B*/*C*), which occurs because a higher-quality individual can expect considerably higher lineage fitness from the usage of a resource than a lower-quality individual can from the same resource.

### 6.2 Genetic bandwagoning can accelerate the evolution of adaptations

By inducing relinquishments of resources by low-quality holders and not doing so in high-quality holders, a bandwagoning variant acts to direct resources from genotypes associated with a long-term fitness disadvantage to genotypes associated with a long-term fitness advantage. The bandwagoning variant becomes selected as its copies “ride on the bandwagon” of the genotypes that offer this long-term fitness advantage. In directing resources to genotypes associated with a long-term fitness advantage, the bandwagoning variant accelerates the fixation of the alleles at other loci that are responsible for this long-term fitness advantage. The acceleration of adaptations that occurs this way is similar to the way that condition-dependent sexual selection can accelerate the fixation of adaptations (Lorch et al., 2003) and also to the way that condition-dependent sex hastens the rate of adaptation by imposing the cost of sex upon low-quality individuals (Hadany and Otto, 2009). In these cases, higher-quality individuals are conferred additional fitness advantages and/or lower-quality individuals disproportionately incur costs, which accelerates adaptive evolution.

## Acknowledgements

The manuscript is dedicated to the memory of Leonard R. Solon (1925 - 2018), World War II veteran, distinguished physicist, and still more memorable grandfather. I further thank Harsh Daga (Vidyalankar Institute of Technology) for writing many of the computer programs used to test the different models of bandwagoning that I have developed over the prior 5 years. Additionally, I thank Josh Mitteldorf for his comments on the manuscript which have substantially improved its clarity. I declare that I have no conflict of interest regarding the publication of this manuscript.

## Appendix

## A1. Calculations of the fitness values of individuals

The fitness lost by a cooperator that foregoes reproduction is equal to the quality of that cooperator. The fitness gained by other individuals from the resources unused by a cooperator that foregoes reproduction is proportional to the fitness lost by that cooperator, but may only be a fraction of this fitness lost because the other individuals in the population may not be able to increase their fecundity enough to garner as much fitness from the resources as the cooperator would have. Therefore, I provide that the fitness obtained by the rest of the population from the fitness foregone by a cooperator is equal to the quality, *Q_i_*, of that cooperator times a factor, *Y*, which represents the fraction of fitness lost by the cooperator that is obtained by the rest of the population. (See Appendix section A3.3 for a discussion about how this way of representing the fitness gained by others is conservative in that it is a simplification that is disadvantageous to the cooperator type.) Since factors limiting reproduction are often resources such as food rather than physiological constraints (Williams, 1966), I assume the value of *Y* is close to unity. Let the quantity of cooperators that have a value of quality below the *P* percentile (and that, therefore, forego reproduction) be denoted *x* and let the sum of the values of quality for these *x* cooperators be denoted *Q^F^*:

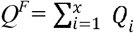

The additional fitness obtainable by the rest of the population from the reproduction foregone by these *x* cooperators is, therefore, *YQ^F^*. The foregone fitness obtainable by other individuals in the population goes to these other individuals in the population according to the assortment (or “relatedness”) in the population and according to the quality of each of the other individuals in the population. This occurs as follows.

In the current model, I follow the custom of treating relatedness, *r*, as an abstract assortment parameter without specifying how this assortment arises in three-dimensional space (Grafen, 1979; Van Cleve and Akcay, 2014; Allen and Nowak, 2015; Cooney et al., 2016; Okasha and Martens, 2016). It is customary to assume that an individual interacts with same-type individuals with probability *r* and with a random individual in the population (possibly same-type and possibly not) with probability 1 - *r*. I assume that if a cooperator foregoes fitness and, therefore, does not use resources it could have, the population is assorted in such a way that *r* percent of the fitness gained by others is obtained by other cooperators and 1 - *r* percent is obtainable by cooperators and defectors. Therefore, fitness of rY*Q^F^* is gained by cooperators and fitness of (1 - r)Y*Q^F^* is gained by all individuals (cooperators and defectors) in the population.

I further assume that an individual obtains the foregone fitness available to individuals of its type in proportion to the individual’s quality. The assumption that higher-quality individuals use a greater proportion of the relinquished resources than other individuals follows from the “good genes” sense of quality as the intrinsic propensity to achieve fitness. Individuals that are of higher quality figure to be advantaged in obtaining resources such as food, territory, and/or mates and in using these resources to gain fitness. Let the sum of quality values of all the defectors in the population be denoted *Q_D_* and let the sum of quality values of all the cooperators with quality at or above the *P* percentile (that is, the cooperators not foregoing reproduction) be denoted *Q_C_*.

Therefore, a cooperator with a quality value of *Q_i_* (that is at or above the *P* percentile) has fitness equal to:

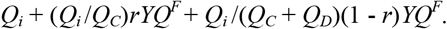

And a defector with a quality value of *Q_i_* has fitness equal to:

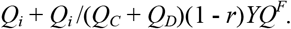

## A2. Guidance for variable values

Literature on humans primarily guides the values used for the simulations because humans are particularly well-studied, so more information is available to inform these variable values. Additionally, human populations are smaller than populations of other well-studied species, which makes the simulations more manageable.

The baseline *N* of 200 approximates Dunbar’s (1993) predicted group size for humans. Human group size probably underestimates human population size. Since a larger population size benefits genetic bandwagoning (see section 3), this baseline number of 200 is conservative with respect to the bandwagoning strategy that is introduced in this manuscript. The baseline value for relatedness is in line with the approximation for human populations calculated by Harpending (2002).

The baseline value for the standard deviation of the distributions from which quality values are selected is 1. The mean quality value across the population is normalized to 10 at the end of each time-step, making the coefficient of variation (Houle, 1992) of quality also equal to 10, which puts it in line with that of the signaling of quality in numerous species (Pomiankowski and Moller, 1995). This standard deviation influences heritability values for reproductive success. The resulting heritability values (Figs. 1, 2 and 3) are of the order of magnitude of values reported empirically for numerous species (Burt, 1995; Merila and Sheldon, 2000).

de Catanzaro’s (1981, 1984) work on depression was mentioned (section 1) as consistent with bandwagoning theory. In the model, cooperators forego reproduction if their quality percentile is below 10, which is in accordance with the percentage of individuals in human populations that battle depression (Kessler and Bromet, 2013).

## A3. Notable simplifications of the model

Simplifications are, by definition, part of models. In this section, I discuss particular simplifications that may be advantageous to either cooperators (that is, simplifications that increase the likelihood of natural selection of genetic bandwagoning) or defectors. These are either simplifications that are traditionally included in Moran models or simplifications included in this manuscript’s particular version of a Moran model in order to model genetic bandwagoning.

In the manuscript’s model, cooperators forego reproduction if their quality is low in order to confer fitness to higher-quality individuals that might be of the same type. The more the lineage fitness gained from such forfeitures exceeds the lineage fitness lost from the forfeitures, the more likely the cooperator type is to be selected. Therefore, simplifications of the model that are advantageous to higher-quality individuals and their lineages (which gain from the forfeitures) are advantageous to the cooperator type, while simplifications that are advantageous to lower-quality individuals and their lineages are advantageous to the defector type. This applies to the simplifications noted in sections A3.1, A3.2, and the first two simplifications noted in section A3.3. Additionally, a simplification of the model that represents the benefit to other individuals from such forfeitures to be higher than otherwise is more favorable to the cooperator type, since assortment in the population is assumed to be positive, so a greater proportion of the benefit goes to other cooperators. This applies to the third simplification noted in section A3.3.

### A3.1 Simplification of uncertain effect: no modeling of sex

If a higher-quality individual and a lower-quality individual were to produce offspring with the same mate, the resulting offspring of the higher-quality individual would have, on average, only one-half the quality advantage over the offspring of the lower-quality individual that the higher-quality parent had over the lower-quality parent. Thus, in a simple model, sex may exert a dilutive influence upon a lineage’s quality advantage over another, which would be a disadvantage to the cooperator type. However, in a more sophisticated model that better resembles dynamics in the wild, there may be assortative mating by quality (Bos et al., 2009; Griggio and Hoi, 2010; Holveck and Riebel, 2010; Holveck et al., 2011; Dakin and Montgomerie, 2014; Veen and Otto, 2015; Schultzhaus et al., 2017), so that higher-quality individuals are more likely to have higher-quality mates. If so, a lineage’s quality advantage over another is more likely to persist, which would benefit the cooperator type. Additionally, if sexual selection were modeled, low-quality individuals that could have reproduced asexually might not be able to find a mate, in which case a low-quality individual would have less to lose by forfeiting resources prior to this anticipated sexual selection, the modeling of which would represent an advantage to the cooperator type.

### A3.2 Simplifications that are advantageous to the cooperator type

1. An individual’s likelihood of being selected to produce offspring is unchanged by whether it has previously been chosen to produce offspring. In the wild, if an individual produces an offspring, it becomes burdened with protecting and providing for that offspring. Consequently, a parent is impaired in the further production of offspring compared to an individual that does not already have offspring. However, in the model, if an individual is chosen to produce the offspring in one time-step, it does not diminish that individual’s likelihood of producing the offspring in a subsequent time-step. This is a simplification that is advantageous to higher-quality individuals, since at any given time-step, they are more likely to have produced prior offspring.
2. There is solely one dimension of quality modeled. In the model, each individual’s quality is characterized solely by a single number. This departs from a wild scenario in which an individual might be, compared to conspecifics, better in one component of fitness (e.g., fighting conspecifics, acquiring territory) and poorer in another (e.g., resisting parasites, evading predators). The modeling of a single dimension of quality is, however, not that different from a wild scenario because an individual’s deficit (or strength) in one component is assumed to influence the individual in another. An individual that is, for example, better at catching prey than another individual is expected, ceteris paribus, to have an advantage in evading predators because it does not have to take as many risks as the other individual in order to catch prey. Likewise, individuals that are better able to avoid wounding or infection are advantaged in their pursuit of prey and their competition with conspecifics, since wounded or infected individuals are impaired in these regards. However, in a more realistic (i.e., wild) scenario, there is the possibility that some traits have a selection differential that is higher than other traits in some time-steps and lower than other traits in other time-steps. For example, if an individual is better than a second individual at capturing prey, while the second individual is better than the first at evading predators, the first individual has more of an advantage when prey are scarce and the second individual has more of an advantage when predators have proliferated. The relevance of more than one trait to an individual’s lineage fitness outlook is suggested by the exhibition of multiple signals, each apparently signaling a particular trait (McCullough and Simmons, 2016). The model does not offer representation to the possibility that there may be genetic variance in multiple facets, with associated selection strengths that differ from one generation to another. This simplification probably benefits the cooperator type because it means that alleles associated with a fitness advantage in one time-step are more likely to be associated with a fitness advantage in subsequent time-steps. However, as noted in Appendix section A2, the baseline parameter values were selected partly so that the heritability values for reproductive success that were generated as the program ran tended to be in line with, or lower than, reported values for multiple species.

### A3.3 Simplifications that are advantageous to the defector type

1. No modeling of differential allocation. In many species, parents have been documented to allocate more resources to offspring produced with higher-quality progenitors (Burley, 1986; Sheldon, 2000; Harris and Uller, 2009; Stiver and Alonzo, 2009; Ratikainen and Kokko, 2010). By investing more resources in offspring from higher-quality mates, parents invest more in offspring that figure to “generate higher returns on investment” than others (Sheldon, 2000; Harris and Uller, 2009). These additional resources to offspring with higher-quality progenitors do not improve the genes of the offspring, but they potentially improve the offspring’s likelihood of surviving to reproductive age and the offspring’s fitness; thereby, they improve the lineage fitness outlook of the higher-quality progenitor. However, in the model, offspring of higher-quality progenitors do not receive an advantage in survival or fitness in addition to their advantage in genetic quality.
2. No modeling of honest signaling of quality. The honest signaling of quality between individuals with competing interests is prevalently reported in the empirical literature on many species (Andersson, 1982; Kirkpatrick, 1982; Andersson, 1994; Hill and Johnson, 2012; Warren et al., 2013). Honest signaling of quality benefits higher-quality individuals because they send honest signals of quality that are more beneficial to them in deciding competitive bouts, obtaining mates, and garnering resources for themselves and their offspring (Kirkpatrick, 1982; Andersson, 1994; Hsu et al., 2006; Rutte et al., 2006). The modeling of honest signaling of quality would benefit the cooperator type due to the benefit it would offer to higher-quality individuals. Yet this benefit is not modeled herein (section 2): In the model, higher-quality individuals do not receive an advantage in survival or fitness in addition to their advantage in genetic quality.
3. The fitness gained by other individuals from the relinquishment of resources by a lower-quality cooperator is proportional to the quality of that lower-quality cooperator. In modeling the evolution of bandwagoning, there must be some manner of representing how a forfeiture of fitness by a lower-quality cooperator winds up increasing the fitness values of other individuals. Bandwagoning theory holds that this occurs as lower-quality individuals do not use resources they could have used for survival and/or reproduction, which allows other individuals to benefit from the usage of these resources. However, in the model, there are no “resources.” Instead, when a lower-quality cooperator loses fitness, the fitness gained by other individuals is proportional to the quality of this lower-quality cooperator. Conceivably, the fitness gained by other individuals could have been set to be proportional to the quality of an average individual in the population. As the low-quality cooperator’s quality is low, setting the fitness gained by others to be proportional to this cooperator’s quality means that the fitness gained by others is represented to be lower than if the fitness gained by others were proportional to the quality of the average individual. A lower quantity of fitness gained by others from the fitness lost by low-quality cooperators is disadvantageous to the cooperator type because assortment in the population is assumed to be positive, so cooperators garner a greater proportion of the fitness gained by others as a consequence of the fitness lost by low-quality cooperators.

## Footnotes

1 The term “quality” is used often but inconsistently in evolutionary literature (Wilson and Nussey, 2010; Bergeron et al., 2011; Hill, 2011). An individual’s genetic quality, i.e., “good genes” is sometimes distinguished from its “phenotypic quality,” i.e.,condition. In the current paper, “quality” is used in the sense of “good genes,” that is, “an individual’s intrinsic propensity or ability to achieve fitness” (Wang et al., 2017) because it has genetic variants that leave it more or less advantaged in prey and territory acquisition or more or less susceptible to, for example, predation or infection.. Variation in genetic quality is assumed to owe to mutations or epimutations having positive-as in the Red Queen’s (Van Valen, 1973; Hartung, 1981; Ridley, 1993; Liow et al., 2011; Brockhurst et al., 2014) and adaptive (Morgan et al., 2012; Waite and Shou, 2012; Hammarlund et al., 2016) races- or negative influences upon quality. An individual’s condition may also be used to indicate its genetic quality (i.e., “good genes”).

2 In some English-speaking countries, a person is said to “jump on the bandwagon” of an entity (for example, another person or a group) if his or her investment in that entity occurs on the condition that it already has a favorable outlook for success and the investment occurs in order to benefit from that impending success. Likewise, a bandwagoning variant determines whether the individual in which it is located is high-quality and if so, it remains latent and if not, it induces the individual to forfeit resources, some of which may be used by individuals with copies of the same bandwagoning variant. Consequently, the net impact is that more resources are spent by high-quality holders. The latent copies of the bandwagoning variant benefit as they hitchhike upon (“ride on the bandwagon” of) high-quality individuals and their lineages.

3 Whether a bandwagoning variant would evolve to induce the relinquishment of some resources or all of the resources its holder could have used can vary in a species-specific and context-specific manner and will depend upon the likelihood that its holder’s quality and lineage fitness outlook change. This is discussed further in section 5.

4 Some of the model’s baseline values are discussed in Appendix section A2 and some simplifications included in the model are discussed in Appendix section A3.

5 This normalization is done so that the mean quality value in the population remains 10 from one generation to another, which keeps the relationship between *Q_σ_* and individuals’ values for quality approximately the same from one generation to another. It also keeps individuals’ values for quality manageable. Otherwise, individuals’ quality values are likely to grow to astronomical levels, since there is, essentially, natural selection for quality: Higher quality values are advantageous in both reproduction and survival and quality is partly heritable.

